# Reversable deformation of artificial cell colony for muscle behavior mimicry triggered by actin polymerization

**DOI:** 10.1101/2021.12.18.473289

**Authors:** Chao Li, Xiangxiang Zhang, Boyu Yang, Feng Wei, Yongshuo Ren, Wei Mu, Xiaojun Han

## Abstract

The mimicry of living tissues from artificial cells is beneficial to understanding the interaction mechanism among cells, as well as holding great potentials in the tissue engineering field. Self-powered artificial cells capable of reversible deformation are developed by encapsulating living mitochondria, actin proteins, and methylcellulose. Upon the addition of pyruvate molecules, the mitochondria produce ATP molecules as energy sources to trigger the polymerization of actin. ATP molecules were produced by mitochondria (2.76×10^10^/ml) with the concentrations of 35.8±3.2 µM, 158.2±19.3 µM and 200.7±20.1 μM by adding pyruvate molecules with the concentration of 3 μM, 12 μM and 21 μM, respectively. The reversible deformation of artificial cells is experienced with spindle shape resulting from the polymerization of actins to form filaments adjacent to the lipid bilayer, subsequently back to spherical shape resulting from the depolymerization of actin filaments upon laser irradiations. The linear colonies composed of these artificial cells exhibit collective contraction and relaxation behavior to mimic muscle tissues. At the stage of maximum contraction, the long axis of each GUV is in parallel to each other. All colonies are synchronized in the contraction phase. The deformation of each GUV in the colonies is influenced by its adjacent GUVs. The muscle-like artificial cell colonies paved the path to develop sustainably self-powered artificial tissues in the field of tissue engineering.

## Introduction

Tissues are comprised of spatially interlinked consortia of cells to exhibit higher-order collective behaviors^1, 2^. The mimicry of living tissues based on the bottom-up fabrication method was beneficial to understanding the interaction mechanism among cells, as well as holding great potentials in the field of bioinspired medical-tissue engineering^3, 4^. The building blocks of artificial tissues include liposomes^5-8^, polymersomes^9, 10^, microdroplets^11-14^, and proteinosomes^15^, etc. The artificial tissues consisted of interlinked proteinosomes with thermoresponsive property exhibited reversible shrinkage due to their thermal effects^15^. The artificial tissues made of lipid protected droplets showed collective deformation under the stimulation of external osmotic pressure^16^. The reported collective deformations of artificial tissues were all triggered by external stimuli. It is a great challenge to trigger the collective deformation of artificial tissue by internal biochemical reactions of the artificial cells.

The artificial cells capable of sustainable energy supply are essential to induce their deformations, consequently to cause the colony deformations. Most of the chemical energy in life is stored in the ATP molecules. ATP molecules were used as energy currency to polymerize actin proteins to form filaments^17, 18^. FoF1-ATPase together with PSII were reconstitute into vesicles to synthesize ATP molecules in the field of artificial cells^19-21^, which involved complicated protein extraction, purification and reconstitution^22^. In living cells, mitochondria are the main factory to produce ATP with the input of pyruvate molecules^23^. There are few reports using mitochondria as power source to produce ATP molecuels^24^.

Muscle tissues are composed of a large number of muscle cells to control body movement and internal organ activity via their contraction functions^25, 26^. The contraction of muscle tissues involves actin filaments and myosin powered by ATP molecules^27^. The reported artificial muscles relied mostly on active materials^28-30^, which included shape memory alloys^31, 32^, liquid crystal elastomers^33, 34^, conductive polymers^35^, carbon nanotubes^36-38^, and hydrogels^39-41^, etc. Hitherto, no research was reported to construct muscle tissue using artificial cells as building blocks.

Herein, we developed an artificial cell capable of reversible deformation as the building blocks to form colonies for mimicking muscle contraction and relaxation behavior. The artificial cell contained mitochondria, actin monomers, and methylcellulose. The pyruvate molecules triggered mitochondria to produce ATP molecules inside the artificial cell, which further initiated the polymerization of actin monomers to form filaments. The artificial cells were deformed by the actin filaments, and recovered by the depolymerization of actin filaments via laser irradiation. Based on the reversible deformation, the linearly hemi-fusion artificial cell colonies were confirmed to exhibit synchronized collective behaviors to mimic muscle tissues. This work paved a path for building self-powered artificial tissues.

## Results and discussion

### Construction of mitochondria-containing GUVs

The mitochondria were extracted from C6 glioma cells. The extracted mitochondria were in rod morphology (Figure 1a) with cross-section diameter of 0.43±0.02 μm and length ranging from 0.70 μm to 1.60 μm. The viability of extracted mitochondria was measured by a dye of JC-1 (5,5′,6,6′-Tetrachloro-1,1′,3,3′-tetraethyl-imidacarbocyanine iodide). The red color indicated the active mitochondria (Supplementary S1). The zeta potential of mitochondria in the buffer solution (pH=8) was -16.6±0.4 mV.

**Figure 1.**
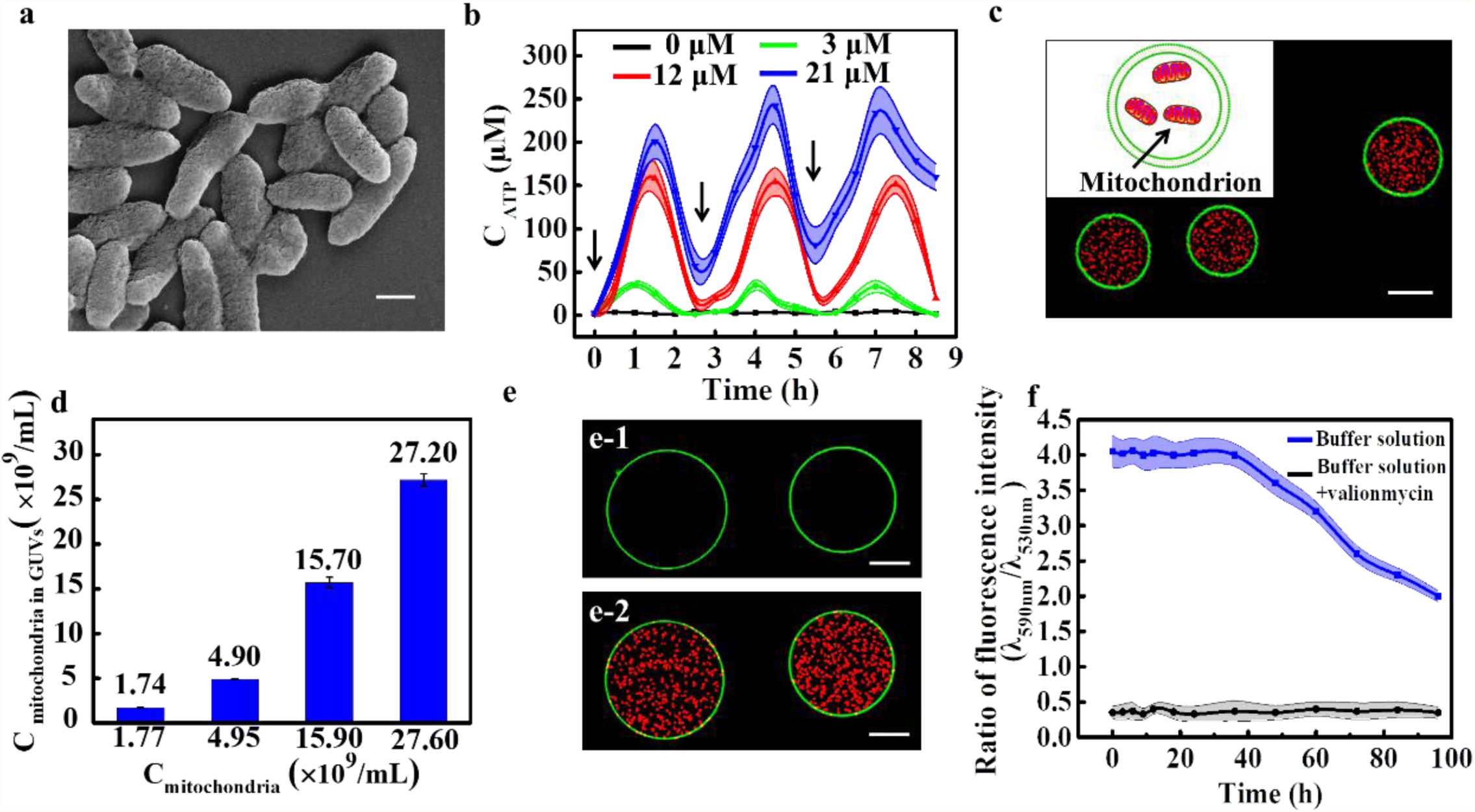
The preparation of mitochondria containing artificial cells. (a) SEM image of extracted mitochondria. (b) Time-dependent ATP generated repeatedly with sequential addition of pyruvate in the buffer solution containing 2.76×10^10^/ml mitochondria (n=3). (c) Fluorescence image of GUVs encapsulated with JC-1 stained mitochondria (red color, 2.76×10^10^ /mL). The inset is the schematic illustration of mitochondria-GUV. The bilayer membrane of GUVs was labelled with NBD-PE (green color). (d) The concentration of mitochondria inside the GUVs as a function of the initial concentration of mitochondria in the buffer solution. (e) Fluorescence images of mitochondria and GUVs before (e-1) and after (e-2) the addition of JC-1 in the solution. (f) Time-dependent viability of mitochondria inside GUVs treated with JC-1 (blue line) and valinomycin (black line). The scale bars were 400 nm in (a), 20 μm in (c) and 10 μm in (e).

Mitochondria produce most of chemical energy stored in ATP molecules for metabolic reactions in cells. Before they were encapsulated into artificial cells for energy supply, the extracted mitochondria were tested for ATP production triggered by pyruvate molecules. The ATP concentration was measured to be 107.8 ±16.3 μM immediately (Supplementary S2) after the mitochondria (2.76×10^10^/ml) were extracted from cells. The inherent ATP molecules were removed by the centrifuging method. The supernatants were replaced with fresh buffer solution eight times to enable the complete removal of free ATP (Supplementary S2). Upon this point, the controllable ATP production of those extracted mitochondria were investigated. It is well known that pyruvate molecules stimulate mitochondria to produce ATP^24^. To prove the ability of mitochondria to regenerate ATP sustainably, pyruvate molecules were added into the mitochondria buffer to trigger the tricarboxylic acid cycle and induce mitochondria to generate ATP (Figure 1b). Upon the addition of 3 μM pyruvate per 2.5 hours into the mitochondria buffer (2.76×10^10^/ml), the concentration of ATP was monitored as a function of time (Figure 1b, green line). The concentration of ATP increased to the peak value of 35.8±3.2 µM in the first 1 hour and decreased to 0 µM after another 1.5 hours. Adding the same concentration of pyruvate again, the extracted mitochondria produced almost the same amount ATP with similar rate (Figure 1b, green line). The ATP generation ability remained at least three repeats. Similarly, the addition of different concentration of pyruvate (12 μM, 21 μM) were investigated. With the addition of 12 μM (Fig. 1b red curve) and 21 μM (Fig. 1b blue curve), the ATP concentration reached 158.2±19.3 µM and 200.7±20.1 μM, respectively. The more addition of pyruvate was, the more ATP concentration gained. They all exhibited the similar patterns (Fig. 1b). For a comparation, there was no significant change of ATP concentration in the control group with no addition of pyruvate molecules (Figure 1b, black line). The physiological activity of mitochondria maintained at least 40 hours after extraction (Supplementary S3). Therefore, the functional mitochondria were successfully extracted, which produced ATP molecules multiple times after repeated addition of pyruvate in a controlled manner. In the following context, the functional mitochondria were used as energy supplier triggered by pyruvate for actin polymerization inside GUVs to form cytoskeletons, and further for cytoskeleton induced GUVs colony deformation to mimic muscle tissue function.

The mitochondria-containing artificial cells (Figure 1c) were prepared using water-in-oil (W/O) emulsion transfer method^42^. The inset of Figure 1c was the cartoon of mitochondria-containing artificial cell. The size distribution indicated that most of GUVs ranging between 20 to 35 µm in diameters (Supplementary S4). In the following experiments, we chose GUVs with diameters between 20 to 35 µm for the observation and statistical analysis. The number of mitochondria encapsulated in the GUVs depended on the initial concentration of the mitochondria during the emulsion formation. The typical confocal fluorescence images of GUVs containing different numbers of mitochondrial were presented in Figure 1c and Supplementary S5. The red dots indicated the active mitochondria due to JC-1 staining. Movie S1 showed the mitochondria moved inside the GUVs, which further confirmed the encapsulation of mitochondria. Take the advantage of emulsion method, almost all the mitochondria were encapsulated in GUVs. The concentrations of mitochondria inside GUVs were almost the same with the mitochondria solution used during the preparation (Figure 1d).

The physiological activity of mitochondria inside GUVs was important for the sustainable energy supply. The viability of mitochondria inside GUVs was monitored using JC-1 staining method. JC-1 dyes were added into the solution after the formation of mitochondria containing GUVs. Figure 1e1 and 1e2 indicated the mitochondria-GUVs before and after the addition of JC-1, respectively. The red dots inside GUVs (Figure e2) confirmed that the JC-1 molecules were able to penetrate the lipid bilayer membrane to stain mitochondria. By measuring the ratio of fluorescence intensity at 590 nm and 530 nm, the mitochondria inside GUVs were confirmed to be alive for 40 hours (Figure 1f, Figure supplementary S6), which was enough for the following experiments.

### Controllable polymerization of actins inside mitochondria-containing-GUVs

Actin filaments are one of the cell cytoskeletons for cell activities^43^. The formation of actin filaments is highly ATP dependent. The mitochondria inside GUVs were the energy power to produce ATP triggered by pyruvate molecules. The distribution of actin filaments inside GUVs was found to be tuned by the actin concentration and the existence of macromolecules (methylcellulose). With lower actin concentration (0.1 mg/mL), the actin filaments were distributed inside the GUVs with short filament morphology, whilst with higher actin concentration (3.3 mg/mL) and methylcellulose, the actin filaments located adjacent to the inner side of lipid bilayer membranes. These two scenarios were investigated respectively.

As illustrated in Figure 2a, mitochondria (2.76×10^10^/ml) and lower concentration of actin monomers (0.1 mg/mL) were encapsulated into GUVs to form the self-energy supplied artificial cells. The other reagents (ADP, Mg^2+^, and Pi) for actin polymerizations were also encapsulated inside the artificial cells. The added pyruvate molecules entered the GUVs from melittin pores in the bilayer membranes, subsequently to trigger mitochondria to produce ATP, which further induced the polymerization of actin monomers into actin filaments to form cytoskeleton (Fig. 2a). When adding 3 μM pyruvate into the buffer every 50 mins, the actins in the GUVs exhibited a stepped polymerization behavior (Figure 2b0-b5), which was confirmed by the fluorescence intensity-time curve (Figure 2c). The actin filaments were stained by TRITC Phalloidin (red florescence) in green GUVs. 3D reconstruction of the GUV in Figure 2b was obtained from serial section images in the Z-stacks taken by a laser confocal microscope^44^ (Movie S2). For a comparation, almost no actin filaments were observed when there were no mitochondria in GUVs (Supplementary S7).

**Figure 2.**
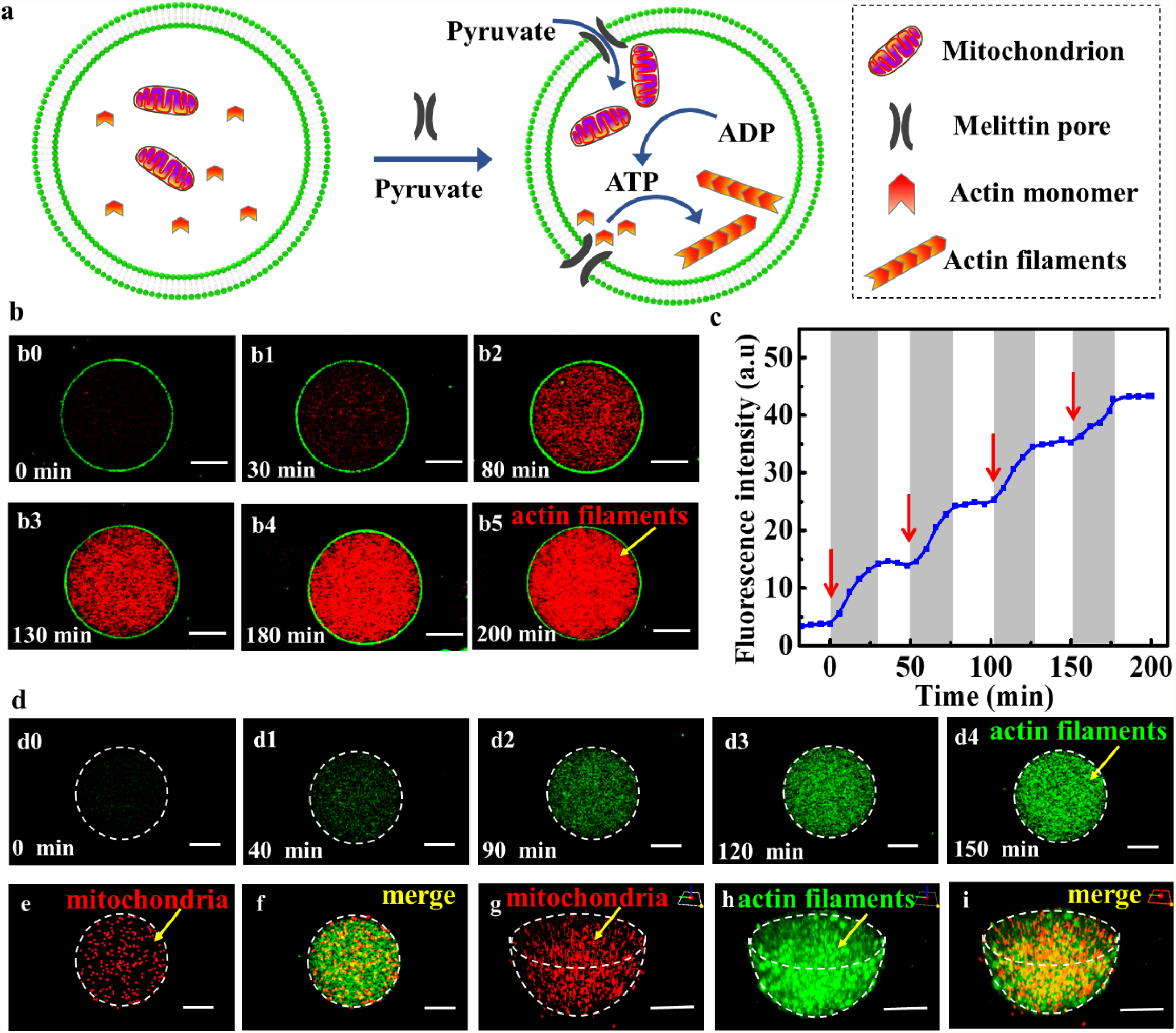
Controllable polymerization of actins. (a) Schematic illustration of actin polymerization in GUVs. (b) Time-dependent fluorescence images of the process of actin polymerization at 0 min (b0), 30 min (b1), 80 min (b2), 130 min (b3), 180 min (b4), 200 min (b5) when adding 3 μM pyruvate into the solution every 50 minutes. (c) The time-dependent average fluorescence intensity of actin filaments in a GUV. (d) Time-dependent fluorescence images of the actin polymerization at 0 min (d0), 40 min (d1), 90 min (d2), 120 min (d3), 150 min (d4) after adding 21 μM pyruvate. (e) Distribution of mitochondria (red fluorescence) in GUVs 150 mins after the actin polymerization. (f) The merged image of (d4) and (e). The three-dimensional reconstruction of mitochondria (g) and cytoskeleton (h) in a GUV after adding 21 μM pyruvate for 150 minutes. (i) The merged image of (g) and (h). The scale bars were 10 µm.

In order to determine the distribution of mitochondria inside GUVs after cytoskeleton formation, the actin filaments and mitochondria were labelled with FITC Phalloidin (green fluorescence) and JC-1 (red fluorescence), respectively. The GUV bilayer was not labelled with fluorescence lipids to avoid the interference. Figure 2d0-d4 showed a process of the actin polymerization with addition of 21 μM pyruvate, where the green color indicated the actin filaments. The mitochondria inside GUVs was evenly distributed after actin polymerization for 150 minutes (Figure 2e). The merging image of the fluorescence images from green channel (actin filaments) and red channel (mitochondria) (Figure 2f) indicated that the mitochondria were evenly mixed with actin filaments. The 3D reconstruction of GUV containing mitochondria and actin filaments in Figure 2g-I exhibited the same results. After the cytoskeleton formation, the mitochondria evenly distributed inside GUVs.

One of the functions of cytoskeleton in cells is to resist external osmotic pressure. The GUVs containing short actin filaments were found to be resistant to the hypertonic conditions (Supplementary S8). Three kinds of GUVs containing different concentrations of actin filaments (0 mg/mL, 0.03 mg/mL, 0.1 mg/mL) were patterned in an acoustic field. Their osmotic pressure resistances were investigated by adding high concentration of glucose outside of GUVs^45, 46^. The GUVs containing more actin filaments exhibited more resistance ability towards the osmotic pressure to maintain GUV spherical morphology.

The interesting results were the reversable deformation of GUVs containing high concentration of actin (3.3 mg/mL), methylcellulose and 2.76×10^10^/ml mitochondria, triggered by the addition of pyruvate molecules (21 μM) (Figure 3a). The GUVs deformed to spindle shape after actin polymerization (middle image of Figure 3a, Figure 3b), which returned spherical shape after depolymerization of actin filaments by laser (540 nm) irradiation (right image of Figure 3a). Figure 3b showed the typical spindle shape of GUVs containing 0.375 w% methylcellulose, 3.3 mg/mL actin, and 2.76×10^10^/mL mitochondria formed after 20 minutes actin polymerization triggered by pyruvate (21 μM). The red regions were the actin filaments stained with red dye of FITC Phalloidin. A 3D image of spindle-GUVs with actin filaments was obtained from serial section images in the Z-stacks taken by a laser confocal microscope (Movie S3), which also confirmed the actin filaments located adjacent to the lipid bilayer due to the excluded-volume effect of methylcellulose^47^. The GUVs were labelled with green NBD PE. We defined a parameter of W-H ratio to describe the deformability, where W and H represented the length of long axis and short axis of spindle GUVs (the inset of Figure 3b). The W-H ratio distribution of the spindle-GUVs (0.375 w% methylcellulose) (Figure 3c) indicated over 65% GUVs with the ratio from 1.2 to 1.7. With mass fraction of methylcellulose to be 0.375%, 80% GUVs were deformed into spindle shape (Figure 3d). With more or less methylcellulose inside GUVs, the deformed GUVs ratio decreased. With more methylcellulose, the solution viscosity increased, which hinder the deformation of GUVs. Therefore, 0.375% was selected as the mass fraction of methylcellulose in subsequent experiments.

**Figure 3.**
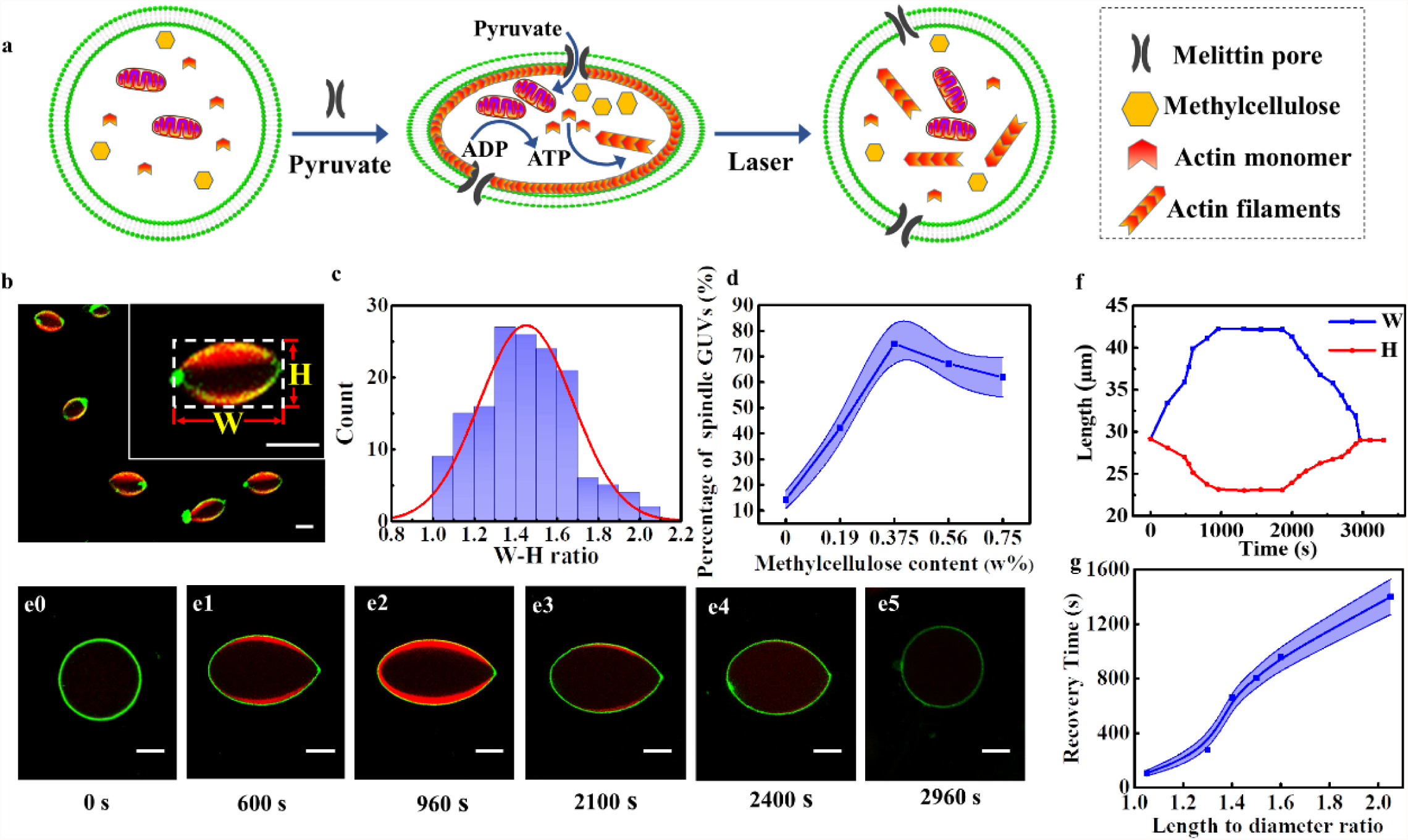
Deformation-reversible GUVs. (a) Schematic of the reversible spherical-spindle-spherical deformation of GUVs induced by polymerization – depolymerization of actin filaments. (b) Typical spindle GUVs containing 0.375 w% methylcellulose, 3.3 mg/mL actin and 2.76×10^10^/mL mitochondria after 20 minutes of actin polymerization triggered by pyruvate (21 μM). (c) The distribution of W-H ratio (the length ratio of long axis over short axis) of spindle-GUVs with methylcellulose (0.375 w%). (d) The percentage of spindle GUVs as a function of mass fraction of methylcellulose in GUVs. (e) Confocal fluorescence images of time-dependent deformation of GUVs at 0 s (e0), 600 s (e1), 960 s (e2), 2400 s (e3), 2700 s (e4) and 2960 s (e5), respectively. (f) Time-dependent W (blue line) and H (red line) values of the GUV in (e) during a reversible deformation process. The scale bars were 10 µm.

The confocal fluorescence images showed the typical deforming process of the GUV from spherical to spindle shape, subsequently back to spherical shape (Figure 3e, Movie S4) upon light irradiation as schematically illustrated in Figure 3a. At 0 s, 21 μM pyruvate was added to the solution to trigger the polymerization of actin inside GUVs. The GUVs at the starting point (Figure 3e0) was in spherical shape. As actin monomer polymerized into red actin filaments, the spherical GUV turned into spindle shape (Figure 3e1). Meanwhile, the W-H ratio of the spindle-shaped GUV increased until it reached the maximum W-H ratio (1.82) at 960 s (Figure 3e2). The lengths of the long axis (W) and the short axis (H) of the GUV were monitored as a function of time (Figure 3f). The laser irradiation caused the actin filaments to gradually depolymerize, which further caused the GUV gradually back to spherical shape (Figure 3e3-e5). Eventually, the spindle-shaped GUV recovered to the spherical shape at 2960 s and maintained spherical shape afterwards (Figure 3e5). In order to determine the distribution of mitochondria after the formation of actin filaments inside GUVs, the actin filaments were stained with FITC Phalloidin (green fluorescence) and mitochondria were stained with JC-1 (red fluorescence) inside the GUV, respectively (Supplementary S9). The GUV bilayer was not labelled with fluorescence lipids to avoid the interference. Supplementary S9 showed the actin filaments (green fluorescence) adhered to the GUV membrane and the mitochondria (red fluorescence) were evenly distributed inside the GUV.

From abovementioned results, the deformation-reversable GUVs were obtained. Given these GUVs have the similar function to muscle fiber cells, GUVs colonies were prepared to mimic muscle tissue contraction and relaxation functions.

### Muscle tissue function mimicry

The artificial cell colony were obtained using an acoustic device equipped with one pair of transducers operating at 6.71 MHz^8, 48^. The GUVs contained 0.375 w% methylcellulose, 3.3 mg/mL actin, and 2.76×10^10^/mL mitochondria. GUVs were trapped in the low acoustic pressure areas (red line areas, Supplementary S10) to generate linear GUVs colonies. The number of GUVs in one colony was adjusted from two to four GUVs by changing the concentration of the GUV (Figure 4a, b and c) ^8^. In order to connect GUVs to each other, 3 mM Ca^2+^ was added into the device to promote hemi-fusion of the adjacent GUVs ^8, 48^. After 20 minutes, the Ca^2+^ solution was replaced with fresh 300 mM sucrose solution containing 4 µg/mL melittin molecules. After another 20 minutes, 21 μM pyruvate was added into the device to trigger the polymerization of actin inside GUVs. Meanwhile, the acoustic field was turned off.

**Figure 4.**
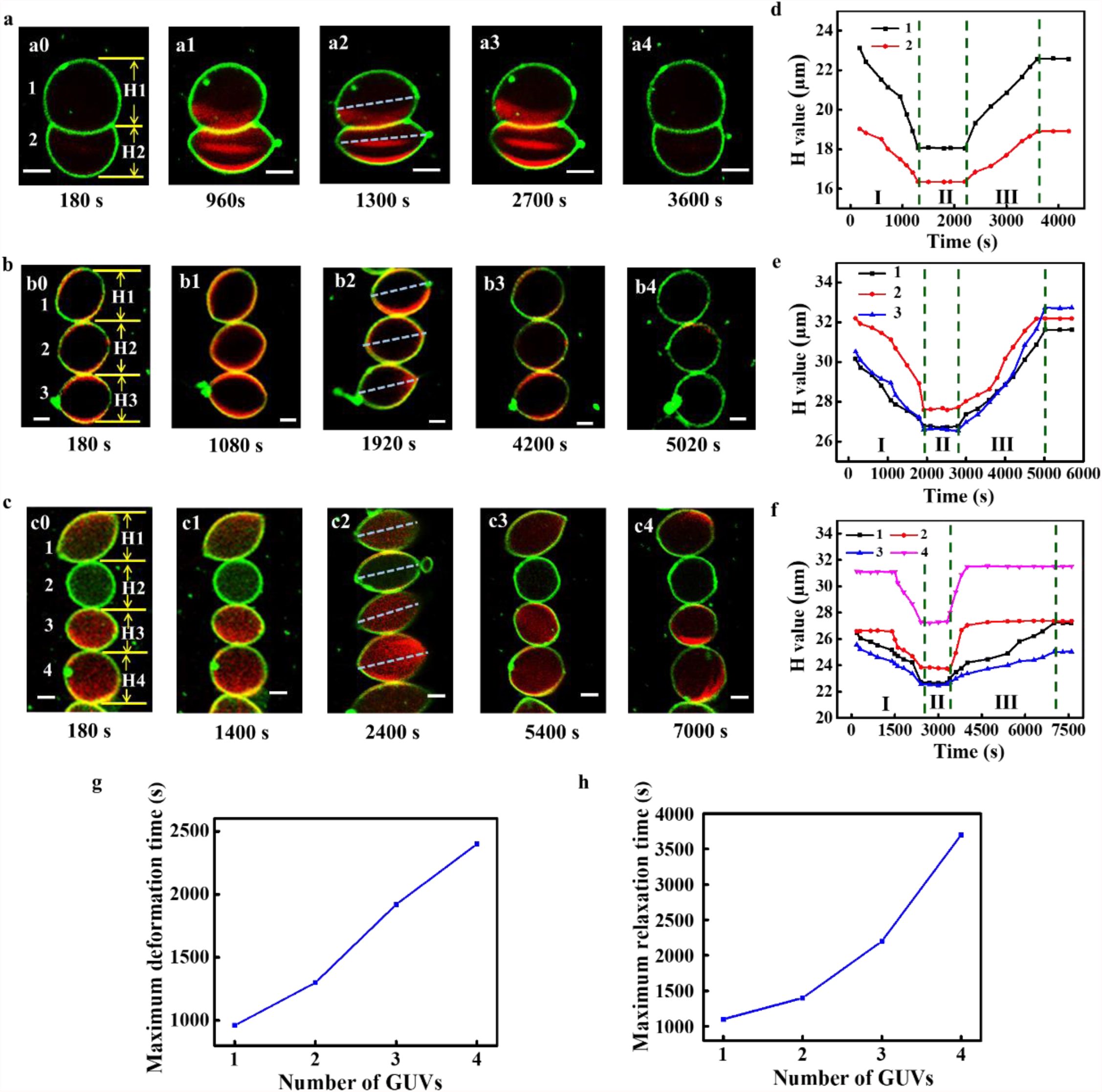
Linear GUVs colonies capable of muscle tissue mimicry. Fluorescence images of time-dependent deformation of two-(a), three-(b), and four-(c) GUV colonies as function of time. The GUVs in (a0) (b0) (c0) were numbered to correspond to the curves in e, f, and g. The dashed lines in (a2) (b2) (c2) represented the long axis of each GUV. (d), e), and f) were the time-dependent H values of each GUV in (a), (b) and (c), respectively. Phase I, II, and III were contraction phase, maintaining phase, and relaxation phase respectively. (g) The maximum deformation time and (h) recovery time of GUV colonies. The scale bars were 10 µm.

Similar to the GUV in Figure 3e, the linear GUVs colonies exhibited reversable collectively deformation. Figure 4a-c and Movie S5-7 showed the process of the deformation of linear colonies containing 2, 3, 4 GUVs, respectively. The H values of all colonies decreased with polymerization of actins inside each GUV, and back to original value upon laser irradiations due to the depolymerization of actin filaments. This behavior was similar to the contraction and relaxation of muscle tissues. The contraction ratio was defined as the maximum change of H value (ΔH) over the original H value (H_0_). The contraction ratio is ∼20.55%, ∼18.35%, ∼12.82%, and ∼12.12% for single GUV, two-GUV colony, three-GUV colony, and four-GUV colony, respectively. The maximum H value decreased against the number of GUVs in the colonies, which indicated the interaction among the GUVs hindered their deformation.

The H values of each GUV in the colonies were monitored as a function of time (Figure 4d-f). All the GUVs deformed from spherical to spindle to spherical shape. The deformation of the GUVs in two-GUV colony was synchronous. The H values of GUV 1 and 2 (Figure 4a) reached the minimum value at the same time (1300 s, Figure4a2, Figure 4d phase Ⅰ), maintained the same values until laser irradiation at 2100 s (Figure 4d, phase Ⅱ), and back to original values at 3600 s (Figure 4a4, Figure 4d phase Ⅲ). This was own to interactions between the hemi-fused two GUVs. The GUVs in three- or four-GUV colonies showed similar synchronous characteristics during the contraction phase (Figure 4f phase Ⅰ, Figure 4g phase Ⅰ).

The time to reach the minimum H values were ∼960 s, ∼1300s, ∼1920s, and ∼2400s for single GUV, two-GUV colony, three-GUV colony, and four-GUV colony, respectively (Figure 4g). The relaxation phases of each colony (Figure 4h) were longer than contraction phase. Interestingly, all GUVs in one colony tended to elongate in the same direction. When the GUVs colonies were in the state of maximum deformation (Figure 4a2, b2, c2), the angles among their long axises (dashed lines in each GUV) were close to 0°, which indicated that each GUV deformed in the same orientation. They all deformed with long axis perpendicular to the colony orientations, i.e., parallel to the semi-fusion areas between adjacent GUVs, which was energy favorite.

All above mentioned results indicated the deformation of each GUV in the colonies was influenced by its adjacent GUVs, consequently resulting the synchronized collective muscle tissue behavior of those colonies.

## Conclusion

A self-powered artificial cell was developed by encapsulating active mitochondria, which was able to produce ATP molecules triggered by pyruvate. The ATP molecules triggered the polymerization of actins inside GUVs. At lower concentration of actin (0.1 mg/mL), short actin filaments were formed and evenly distributed inside GUVs. At higher concentration (3.3 mg/mL) and together with methylcellulose, the actin filaments were located adjacent to the inner lipid bilayer of GUVs to induce the deformation of GUVs into spindle shape. Upon the laser irradiation, the spindle GUVs were back to spherical shape. Thus, the artificial cells capable of reversible deformation were demonstrated. Furthermore, the self-powered linear artificial cell colonies were fabricated using those artificial cells to mimic the contraction and relaxation of muscle tissues. The maximum deformation ratio H values decreased in an order of two-GUV colony, three-GUV colony and four-GUV colony. The contraction of those colonies exhibited synchronized collective behavior due to the interaction among GUVs. The muscle-like artificial cell colonies paved the path to develop more complicated artificial tissues both in fundamental and practice point of views.

## Experimental

### Materials

1-palmitoyl-2-oleoyl-glycero-3-phosphocholine (POPC) and cholesterol (Chol) were purchased from Avanti Polar Lipids (USA). 1,2-dioleoyl-sn-glycero-3-phosphoethanolamine-N-(7-nitro-2-1,3-benzoxadiazol-4-yl) (NBD PE) was obtained from Invitrogen (China). Cell Mitochondria Isolation Kit, (hydroxymethyl) aminomethane hydrochloride (Tris-HCl) (pH=8.8) and DL-Dithiothreitol (DTT) were purchased from Beyotime (China). ATP Determination Kit was purchased from Nanjing Jiancheng Bioengineering Institute (China). Actins were purchased from Cytoskeleton Inc. (USA). Mitochondrial Membrane Potential Assay Kit with JC-1 and TRITC phalloidin, and FITC phalloidin were purchased from Solarbio (China). Pyruvate was purchased from Aladdin (China). Sucrose, glucose, calcium chloride (CaCl_2_), magnesium chloride (MgCl_2_), sodium dihydrogen phosphate (NaH_2_PO_4_), potassium phosphate monobasic (KH_2_PO_4_), adenosine diphosphate (ADP), methylcellulose and melittin (85% HPLC) were purchased from Sigma (China). High glucose DMEM, phosphate buffer saline (PBS), and trypsin-EDTA were purchased from Corning (USA). Fetal bovine serum (FBS) was purchased from Gibco (USA). Penicillin-Streptomycin Solution (100X) was purchased from Beyotime (China). Millipore Milli-Q water with a resistivity of 18.2 MΩ cm was used in the experiments.

### C6 glioma cells cultures

C6 glioma cells were cultured in DMEM medium (High Glucose DMEM, 10% fetal bovine serum, penicillin (100 units/mL), and streptomycin (100 µg/mL)) at 37 °C and 5% CO_2_ atmosphere.

### Mitochondria extraction

Mitochondria were extracted following the protocol of the Mitochondria Isolation Kit. In brief, C6 cells were firstly broken by the homogenizer in lysis buffer solution. The resultant solutions were centrifuged (200 g, 10 minutes) at 4°C, followed by centrifuging the supernatant (12000 g, 10 minutes) to collect the mitochondria in the precipitates. Afterwards, the extracted mitochondria were resuspended in buffer solution for future use.

### ATP measurements

The ATP-producing capacity of extracted mitochondria was investigated in the buffer solution containing 300 mM sucrose, 5 mM Tris-HCl (pH=8), 5 mM ADP, 0.5 mM DTT, 0.2 mM CaCl_2_, 2 mM MgCl_2_, 3 mM NaH_2_PO_4_ and 3 mM KH_2_PO_4_. ATP Determination Kits were used to detect ATP produced by mitochondria. Creatine kinase catalyzes adenosine triphosphate and creatine to generate creatine phosphate, which is detected by the phosphomolybdic acid colorimetric method.

### Measurement of mitochondria concentration

1 µL mitochondria suspension solution was diluted 100 times, which was subsequently dropped onto the cover slide in the confined regions. All mitochondria in the solution sedimented onto the surface of the cover slide after 60 minutes, which were then counted using a confocal fluorescence microscope. Fluorescence microscopy images were recorded in different area on the cover slide. The mitochondria were evenly distributed on the slide surface with a suitable density. The mitochondria concentration (c) was calculated from 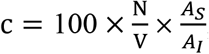, where N represented the average number of the mitochondria in the images (n=5), V was the volume (1µL) of added mitochondria suspension solution, *A*_*S*_ is the total area of the confined region of the cover slide, and *A*_*I*_ was the area of each image. Thus, we obtained the mitochondria concentration in a form of number per volume.

### Preparation of mitochondria-containing GUVs

Mitochondria-containing GUVs were prepared by emulsion transfer method. In brief, 0.5 mg POPC, cholesterol, and NBD PE (65: 30: 5, w/w/w) were dissolved in 50 μl chloroform. 500 µl liquid paraffin was added to the above chloroform solution. The mixed solution was incubated at 80°C for 30 minutes to vaporize chloroform and then cooled down to room temperature to obtain the lipid oil phase solution. The aqueous solution was prepared with mitochondria (1.77×10^9^/ml, 4.95×10^9^/ml, 1.59×10^10^ /ml, and 2.76×10^10^/mL), 300 mM sucrose, 5 mM Tris-HCl (pH= 8), 0.1 mg/mL actin, 5 mM ADP, 0.5 mM DTT, 0.2 mM CaCl_2_, 2 mM MgCl_2_, 3 mM NaH_2_PO_4_, 3 mM KH_2_PO_4_, 0.1 μM TRITC phalloidin (or FITC phalloidin) and methylcellulose (if necessary). 300 µl of the lipid oil phase solution and 30 µl aqueous solution were mixed by vortexing for 30 s to generate water-in-oil emulsion solution. The emulsion solution was added gently into 200 µL isotonic glucose solution in a 1.5 mL centrifuge tube, followed by centrifuging (10000 g, 30 minutes) to generate giant unilamellar vesicles (GUVs) in the precipitates. Thus mitochondria-containing GUVs were obtained. The concentration of mitochondria inside GUVs was determined by the difference between the initial and non-encapsulated mitochondria concentration. The concentration of non-encapsulated mitochondria was obtained by measuring the supernatant of centrifuged just-prepared mitochondria-containing GUVs. The GUVs containing 2.76×10^10^/mL mitochondria were used in all the experiments. Melittin molecules (4 µg/mL) were used to form the melittin pores in the bilayer membrane of GUVs to allow the penetration of pyruvates.

### Mitochondrial viability assay

The inner mitochondrial membrane (IMM) potential (Δψm) indicates the electrochemical gradient across the IMM. Δψm serves as an indicator of mitochondrial activity and the capacity to generate ATP by oxidative phosphorylation. The Δψm was tested using Mitochondrial Membrane Potential Assay Kit with JC-1. Mitochondria with high viability showed red fluorescence (λ_ex_/λ_em_ = 490/590 nm) in the mitochondrial matrix. On the contrary, the solution showed green fluorescence (λ_ex_/λ_em_ = 490/530 nm). 0.9 mL JC-1 was added into 0.1 mL mitochondria solution (2.7×10^10^/mL) for 10 min after the mitochondria were extracted. The viability of the mitochondria was assayed by monitoring the ratio of red/green fluorescence at 0, 3, 6, 9, 12, 18, 24, 36, 48, 60, 72, 84 and 96 h. As a control, mitochondria were pretreated by the buffer solution containing 10 μM valinomycin, which dissipated the Δψm.

### Formation of the actin filaments inside GUVs

The GUVs containing 2.7×10^10^/mL mitochondria and 0.1 mg/mL actin monomer were treated with 4 µg/mL melittin for 20 mins to form melittin pores in the membranes, which allowed the penetration of pyruvates. For stepped polymerization of actin inside GUVs, 3 µM pyruvate was added into the mitochondria-containing GUVs solution every 50 mins to trigger the polymerization of actins to form short filaments inside GUVs.

When the GUVs contained 7×10^10^/mL mitochondria, 3.3 mg/mL actin monomer, and certain amount of methylcellulose, the actin filaments were formed adjacent to the lipid bilayer upon the addition of 21 µM pyruvate. Consequently, the GUVs deformed into spindle shape, which were back to sphere shape upon laser (540 nm) irradiations.

### The reversable deformation of GUVs colonies

GUVs capable of reversible deformation was added into an acoustic device equipped with one pair of transducers operating at 6.71 MHz. The solution was replaced by 300 mM sucrose containing 3 mM Ca^2+^ for 20 mins after the GUVs were patterned into linear colonies. Afterwards, the solution was replaced by fresh 300 mM sucrose solution containing 4 µg/mL melittin for 20 mins, following by adding 21 µM pyruvate into the solution to cause the deformation of GUVs colonies. The recovery of deformed GUV colonies was realized by laser (540 nm) irradiation. The time of deformation and recovery were recorded.

### Instruments

The morphology of mitochondria was characterized by scanning electron microscopy (Quanta 200 FEG, Netherlands). The laser confocal microscope (Olympus FV 3000, Japan) was used to obtain all fluorescence images. The acoustic field was generated by a pair of piezoelectric transducers (PZT, Noliac, NCE 51, Denmark) and a signal generator (Agilent 33220a-001, USA).

## Acknowledgements

This work was supported by the National Natural Science Foundation of China (Grant No. 22174031, 21929401, 22111540252, 21773050), the Fundamental Research Funds for the Central Universities (HIT.OCEF.2021026), and the Heilongjiang Touyan Team (HITTY-20190034).

## Author contributions

X.J.H. supervised the research. X.J.H., C.L., and X.X.Z. conceived and designed the experiments. C.L. and X.X.Z. performed experiments. X.J.H., C.L., X.X.Z., B.Y.Y., F.W., Y.S.R and W.M. analyzed the data. X.J.H., C.L. and X.X.Z. wrote the paper, and all authors commented on the paper. C.L. and X.X.Z. contributed equally to this work.

