## Supporting information for "Reversable deformation of artificial cell colony for muscle behavior mimicry triggered by actin polymerization"

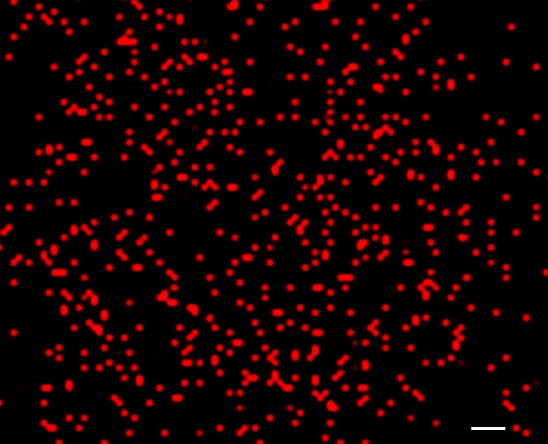

**Supplementary S1.** A fluorescence image of JC-1 labeled mitochondria (red color). The scale bar was 5 μm.

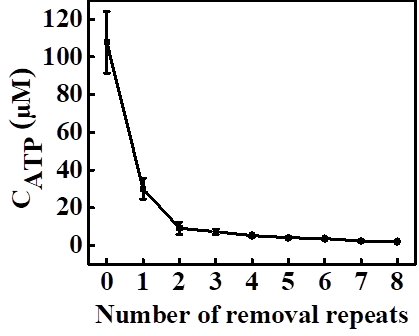

**Supplementary S2.** Concentrations of ATP in the supernatants of the buffer solution containing 2.76×10^10^/ml mitochondria as a function of removal repeats.

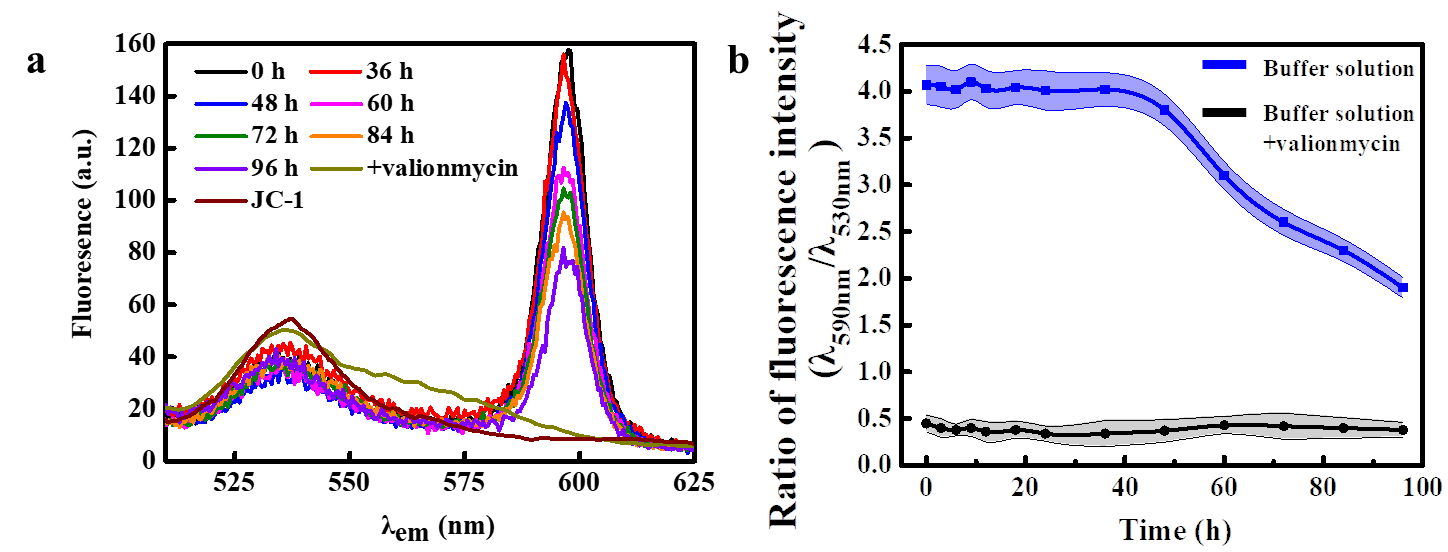

**Supplementary S3.** The physiological activity of mitochondria at different time in buffer solution. (a) Fluorescence spectra of mitochondria in buffer solution treated with JC-1 and with valinomycin after extraction at different time. (b) Time-dependent viability of mitochondria in buffer solution treated with JC-1 (blue line) and with valinomycin (black line).

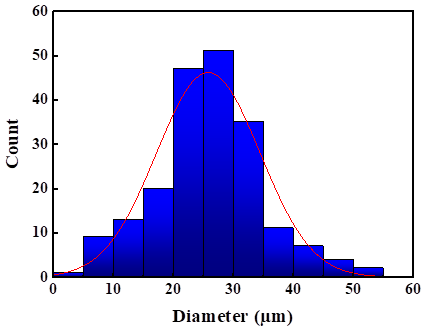

**Supplementary S4.** Diameter distribution of mitochondria-containing GUVs.

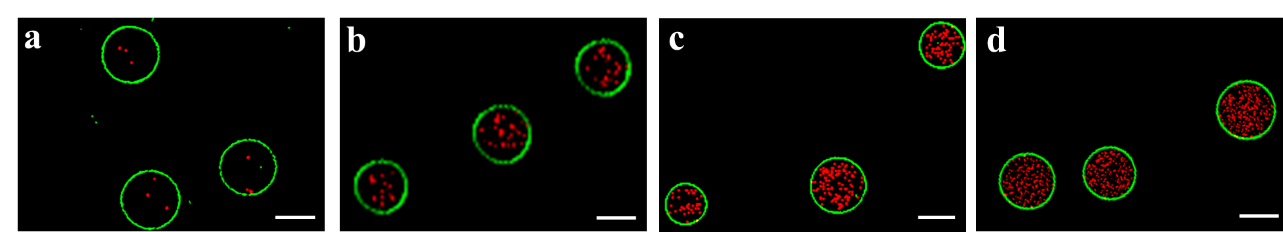

**Supplementary S5.** Fluorescence images of mitochondria-containing artificial cells with mitochondria concentrations of 1.77×10^9^ /ml (a), 4.95×10^9^ /ml (b), 1.59×10^10^ /ml (c), 2.76×10^10^ /mL (d). The scale bars were 20 μm.

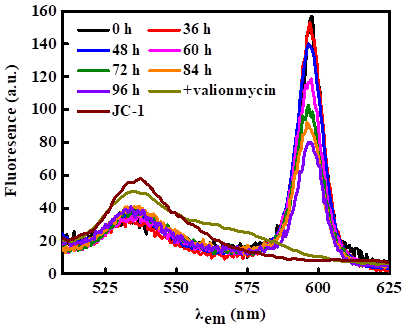

**Supplementary S6.** Fluorescence spectra of mitochondria in GUVs treated with JC-1 and with valinomycin after extraction at different time.

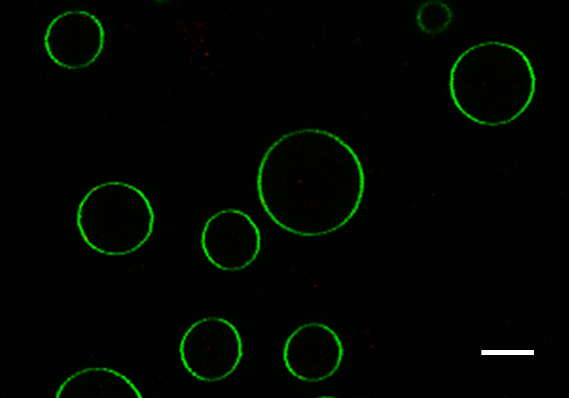

**Supplementary S7.** Fluorescence image of the green fluorescence GUVs with melittin pores containing 0.1 mg/mL actin without mitochondria. When pyruvate (21 μM) was added, no actin filaments (red fluorescence) was observed. The scale bar was 20 µm.

**Osmotic stability of GUV containing actin filaments**

Three kinds of GUVs containing different concentrations of actin filaments (0 mg/mL, 0.03 mg/mL, 0.1 mg/mL) were prepared and patterned in low pressure nods in the acoustic field to generate triangular micro-arrays, respectively (Supplementary S8). The osmotic pressure was adjusted by varying the concentrations of inside sucrose and outside glucose of GUVs. The inside sucrose concentration was 300 mM. GUVs without actin filaments started to deform into vesicle-in-vesicle (VIV) structures under 30 mM (0.73 atm) osmotic pressure (Supplementary S8a3). GUVs with 0.03 mg/mL actin filaments started to deform into VIV structures under 100 mM (2.45 atm) osmotic pressure (Supplementary S8b3). GUVs with 0.1 mg/mL actin filaments started to deform into VIV structures under 300 mM (7.34 atm) osmotic pressure (Supplementary S8c3). The more actin filaments were inside GUVs, the more osmotic pressure was needed to deform into VIV structures.

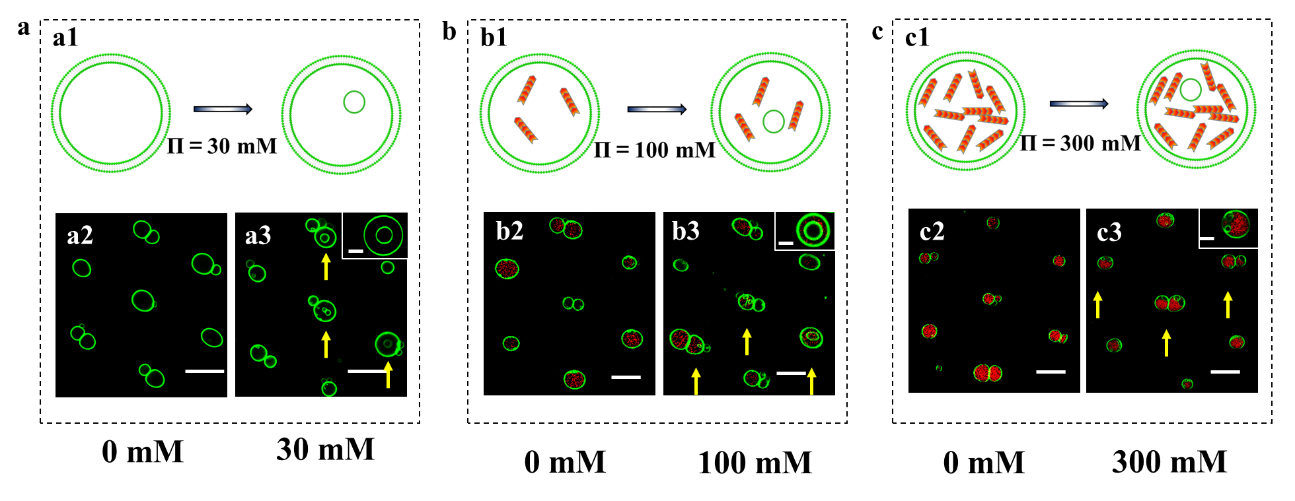

**Supplementary S8. The osmotic pressure resistant ability of GUVs with actin filaments.** Schematic illustration of GUVs containing 0 (a1), 0.03 mg/mL (b1), and 0.1 mg/mL (c1) actin filaments before and after treating with osmotic pressure. Confocal fluorescence images of GUVs containing 0 (a1), 0.03 mg/mL (b1), and 0.1 mg/mL (c1) actin filaments before (a2, b2, c2, respectively) and after (a3, b3, c3, respectively) treating with osmotic pressure. The scale bars were 50 μm.

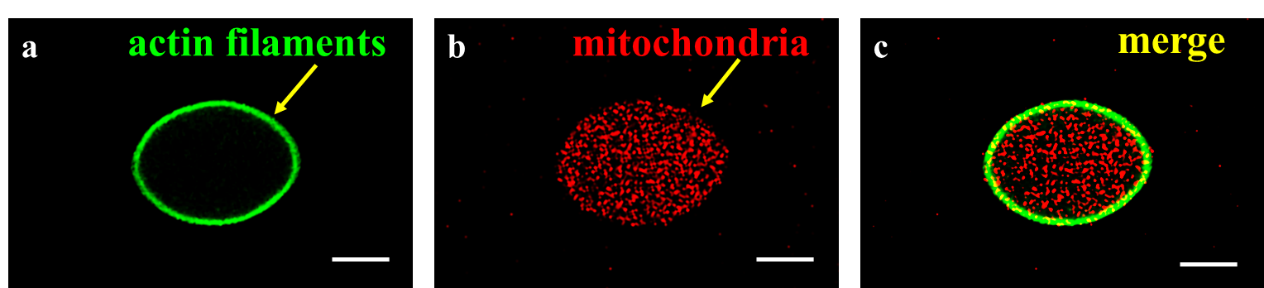

**Supplementary S9.** (a) Fluorescence image of the actin bundles (green) adjacent to the lipid bilayer of GUV. (b) Fluorescence image of mitochondria (red) inside the GUV. (c) The merged image of (a) and (b). The scale bars were 10 μm.

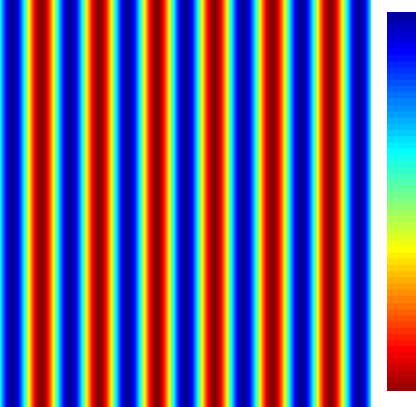

**Supplementary S10.** Simulation of the acoustic pressure distribution in the acoustic field generation device equipped with one pair of transducers operating at 6.71 MHz. Red areas indicated the acoustic low pressure areas, while the blue areas indicated the high pressure areas.
